# Opposing BOLD signals and oxygen metabolism largely arise from statistical uncertainty in metabolic estimates

**DOI:** 10.64898/2026.04.21.719913

**Authors:** Ole Goltermann, Alexander Huth, Christian Büchel

## Abstract

Recent work by Epp et al. (2025) reported widespread voxel-wise sign discordance between task-evoked blood-oxygenation-level-dependent (BOLD) responses and estimated changes in cerebral metabolic rate of oxygen (ΔCMRO_2_), raising important questions about the interpretability of BOLD functional magnetic resonance imaging. Reanalysing the dataset, we found that ΔCMRO_2_ estimates showed substantial voxel-wise variability across participants, consistent with the noise sensitivity of model-based metabolic estimates. When this variability was taken into account, 77.2% of voxels could not be robustly classified, as ΔCMRO_2_ effects lacked sufficient statistical support to determine concordance or discordance. Where classification was possible, positive BOLD responses were predominantly concordant with metabolism, whereas discordance was considerably higher for negative BOLD responses. These findings suggest that the observed BOLD–metabolism discordance reported previously largely reflects statistical uncertainty in CMRO_2_ estimates rather than widespread physiological sign reversal.

## Introduction

In a recently published study, Epp et al. (2025) report widespread voxel-wise discordance between functional magnetic resonance imaging (fMRI) based blood-oxygenation-level-dependent (BOLD) signal changes and quantitative estimates of cerebral oxygen metabolism, concluding that BOLD responses frequently reflect metabolic changes of the opposite sign to those predicted by canonical neurovascular coupling. The findings have been widely covered by news outlets and have prompted broad discussion within the field about the physiological interpretability of BOLD fMRI and its implications for prior and ongoing neuroimaging work.

Extensive empirical work has examined how BOLD signal changes relate to underlying neuronal and metabolic processes. Simultaneous electrophysiology-fMRI experiments in non-human primates demonstrated that BOLD amplitudes covary with local field potentials (Logothetis et al., 2001), and gamma-band activity was shown to be linked to local hemodynamic signals and to predict BOLD signal variance (Magri et al., 2012; Niessing et al., 2005). However, it is also established that the origin and interpretability of BOLD signal is complex, dynamic and far from understood (see for reviews: Buxton, 2012; Kim & Ogawa, 2012; Logothetis, 2008). In particular, negative BOLD responses remain difficult to interpret, as signal reductions can have heterogeneous metabolic underpinning (Godbersen et al., 2023; Stiernman et al., 2021). Whereas most of earlier work emphasized the need for caution in interpreting BOLD signal changes, Epp et al. suggest that the implications may be more substantial, reporting that the canonical interpretation of BOLD signal changes may lead to misinterpretation in approximately 40% of reported BOLD responses.

Epp et al. defined responses as *discordant* when they found opposing signs between task-evoked changes in BOLD (ΔBOLD) and quantitative cerebral metabolic rate of oxygen (ΔCMRO_2_). ΔCMRO_2_ was estimated using a biophysical model of oxygen delivery and extraction (He & Yablonskiy, 2007). The model combines arterial spin labeling–based cerebral blood flow (CBF) with venous deoxyhaemoglobin estimates derived from BOLD and R2′ relaxometry to infer changes in oxygen consumption. This approach provides an indirect fMRI-based alternative to oxygen-tracer positron emission tomography (PET), which is considered the gold standard for CMRO_2_ measurement (Ito et al., 2021). As with other quantitative fMRI approaches, ΔCMRO_2_ estimates depend on multiple model assumptions and are inherently noisier than PET-based metabolic measures (see for example Bright et al., 2019). Following this, one alternative possibility raised by Huth (2026) is that the reported discordance is due to measurement noise. Using simulations in which effect sizes were taken from the study and noise levels were set to reasonable, moderate values, Huth showed that even perfectly aligned underlying signals yield discordance rates of about ∼40%, demonstrating that the observed discordance could arise as a consequence of noisy estimates rather than opposing physiology. To our surprise, Epp et al. did not explicitly account for noise or variability in general in CMRO_2_ estimates, instead treating group-averaged values as a direct marker for metabolic changes, no matter how much single participant estimates varied. Building on this observation, and enabled by the open and transparent sharing of code and data by Epp et al., we reanalysed the original dataset to assess to what extent voxel-wise concordance or discordance classifications were supported by statistically robust ΔCMRO_2_ estimates.

## Results and Discussion

The dataset comprises quantitative fMRI measurements of hemodynamic and metabolic parameters, including BOLD signal, cerebral blood flow (CBF), cerebral blood volume (CBV), effective transverse relaxation time (T2*), reversible transverse relaxation rate (R2′), and model-based estimates of cerebral metabolic rate of oxygen (CMRO_2_). Data were acquired under two task conditions (calculation task and control, see Methods), and task-evoked changes were quantified as percent signal differences between conditions (ΔBOLD and ΔCMRO_2_). We first replicated the original concordance analysis by calculating voxel-wise sign mismatch between ΔBOLD and ΔCMRO_2_. As in Epp et al., we identified voxels showing statistically significant task-evoked BOLD responses (Fig. 2A; hereafter ‘BOLD activation mask’) and compared the signs of group-averaged ΔBOLD and ΔCMRO_2_ within this mask. Consistent with Epp et al., approximately 35% of voxels exhibited opposing signs of ΔBOLD and ΔCMRO_2_. However, this analysis assumes reliable and participant-consistent voxel-wise ΔCMRO_2_ estimates without formally evaluating either. Two examples illustrate this issue.

### Example 1 (high variance)

A voxel shows a significant ΔBOLD of +0.5%. The group-mean ΔCMRO_2_ is −5%, and it is therefore labelled discordant. However, participant-level ΔCMRO_2_ estimates range from −50% to +40% with a high standard deviation. Such variability indicates that the group mean might not be a robust measure for the central tendency of the data in this case. Despite this, the voxel would still be classified as discordant.

### Example 2 (inconsistent sign across participants)

Another voxel shows ΔBOLD of +0.5% and a small positive group-mean ΔCMRO_2_ of +0.2%, and is therefore labelled concordant. Yet 22 out of 40 participants show opposite signs of ΔBOLD and ΔCMRO_2_ (concordance rate = 0.45). Despite this discordant pattern on participant-level, the voxel is classified as concordant.

In both cases, classification is driven solely by the group mean, without evaluating whether the direction of ΔCMRO_2_ is statistically robust. Importantly, replacing the mean with the median (as done by Epp et al.) does not solve this issue, as neither statistic captures uncertainty or sign consistency. To quantify the issue of variability in ΔCMRO_2_ across participants, we examined variance descriptively across contrasts (Fig. 1). For each measure of interest (CBF, CBV, T2*, R2’, and CMRO_2_), we computed the coefficient of variation (CV) in each voxel as the standard deviation of the measure across subjects divided by the mean across subjects. This ratio indicates how consistent each measure is across participants, reflecting both inter-individual variability and noise. In both baseline and task conditions, CMRO_2_ showed the highest CV. In percent signal change maps, BOLD exhibited a left-shifted and markedly narrower CV distribution, indicating lower relative variability across participants, whereas CMRO_2_ showed both higher central CV values and substantially broader spread. Quantitatively, the median CV of ΔCMRO_2_ [= 4.72] was 26.1% higher than ΔBOLD [=3.74] across all cortical voxels and 127.8% [3.72 against 1.63] higher within BOLD-significant voxels, indicating markedly elevated inter-individual variability of metabolic estimates.

**Figure 1.**
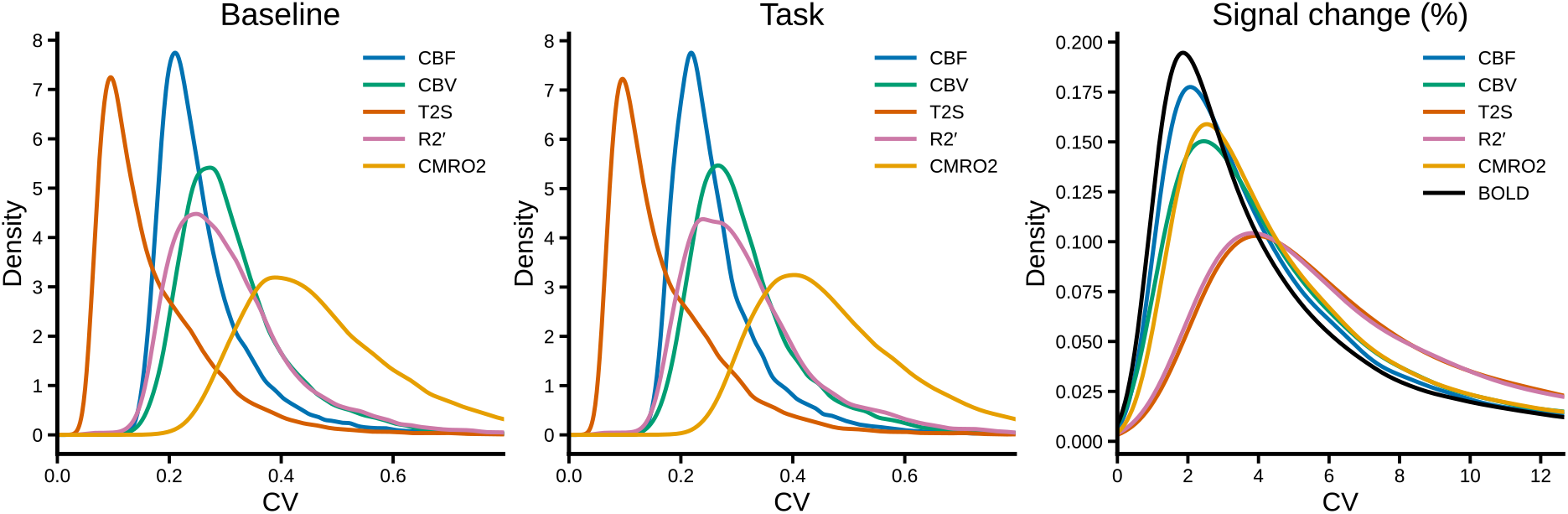
Kernel density distributions of voxel-wise coefficients of variation (CV = SD/|mean| across participants) for CBF, CBV, T2*, R2′, CMRO_2_, and BOLD. Left and middle panels show CV distributions under baseline [control task] and task [calculation] conditions, respectively. Right panel shows CV distributions of percent signal change (PSC) estimates.

These observations demonstrate that ΔCMRO_2_ exhibits substantial variance, raising concerns about sign classification based solely on group averages. To address this limitation, we implemented two complementary inferential approaches. The first evaluated the stability of the sign relationship between ΔBOLD and ΔCMRO_2_ across participants. The second tested whether ΔCMRO_2_ differed significantly from zero and assigned concordant/discordant labels only for voxels showing a significant metabolic effect; all other voxels were categorized as showing no significant ΔCMRO_2_ change.

First, we tested whether the voxel-wise sign concordance rate across participants differed from chance. For each voxel, we computed the proportion of participants showing matching signs (positive or negative) for ΔBOLD and ΔCMRO_2_, and tested this proportion using a binomial test against a null hypothesis of 0.5, corresponding to chance-level agreement between the signs of ΔBOLD and ΔCMRO_2_. Voxels with concordance rates not significantly different from 0.5 cannot be robustly classified as either concordant or discordant. Across voxels, concordance rates clustered near 0.5 (mean = 0.54; Fig. 2B). Only 65 out of the 19,190 voxels in the BOLD activation mask (0.3%) were statistically classified as concordant and only 10 (0.05%) as discordant (FDR-corrected q = 0.05; empirical thresholds: p_concordant_ ≥ 0.81, p_discordant_ ≤ 0.19). Although this test is conservative, as it requires a strong and statistically reliable deviation from chance across participants, this analysis indicates that, for most voxels, the sign relationship between ΔBOLD and ΔCMRO_2_ is not consistent across participants and group-based classification should be interpreted with caution.

**Figure 2.**
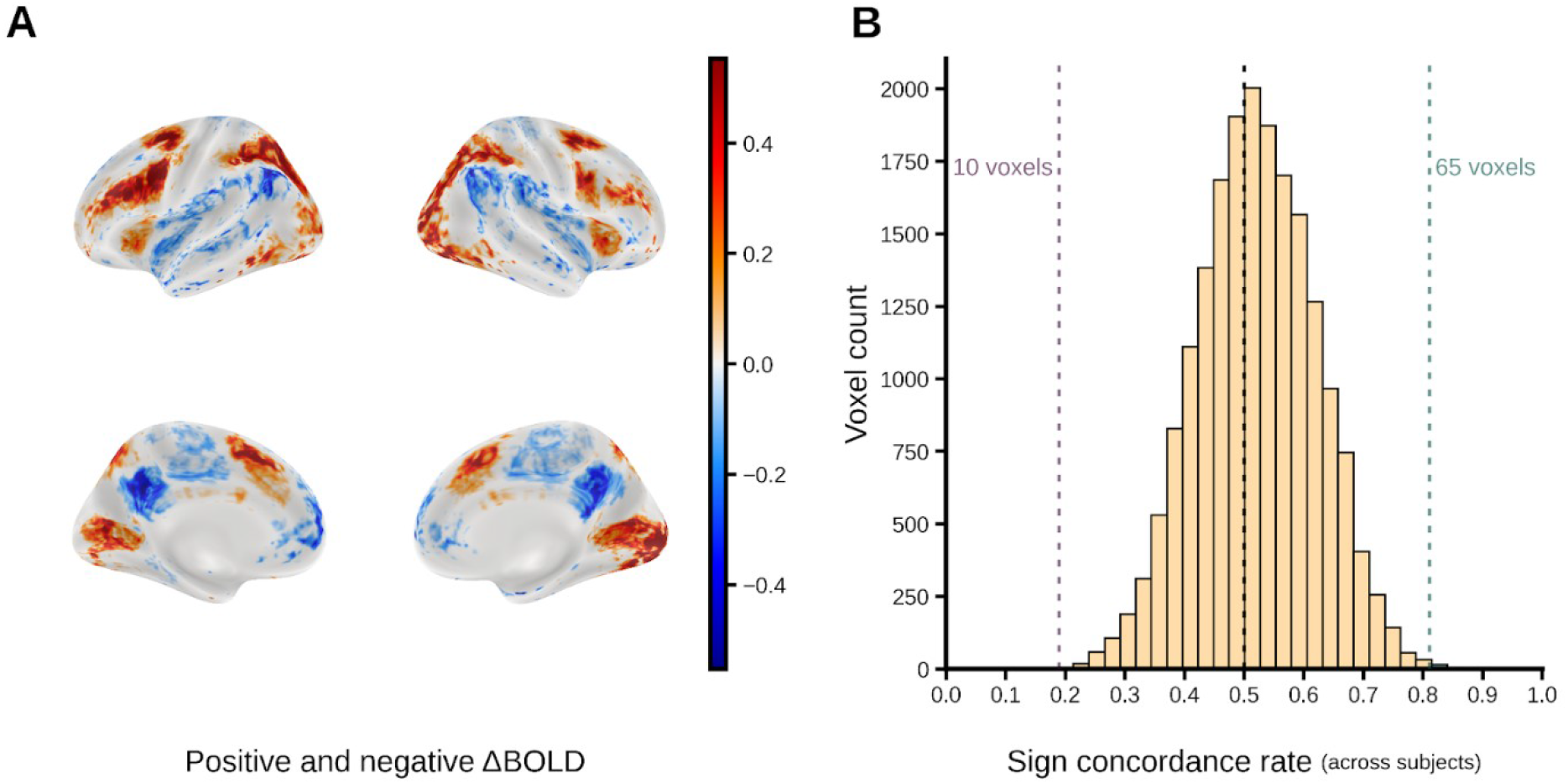
**A** Cortical surface maps showing the mean change in BOLD signal (ΔBOLD) across participants for all voxels included in the BOLD activation mask. Warm colors (red–yellow) indicate positive ΔBOLD values, whereas cool colors (blue) indicate negative ΔBOLD values. The color bar represents mean ΔBOLD (percent signal change). **B** Histogram of voxel-wise sign concordance rates across participants within the BOLD activation mask. The concordance rate reflects the proportion of participants showing the same sign (positive or negative) of ΔBOLD and ΔCMRO_2_ at a given voxel. Vertical dashed lines denote q(FDR)<0.05 significance thresholds for classifying voxels as discordant (sign concordance below chance) or concordant (sign concordance above chance). The numbers above the dashed lines indicate the number of voxels falling beyond each threshold.

Second, we applied a group-level statistical criterion that likely provides a less conservative assessment of sign relationships than the participant-level concordance test described above. Specifically, we tested whether ΔCMRO_2_ differed significantly from zero across participants and classified voxels as concordant (matching signs of significant ΔBOLD and significant ΔCMRO_2_) or discordant (opposing signs) when ΔCMRO_2_ reached statistical significance at the group level, and otherwise as showing no significant ΔCMRO_2_ effect (see Methods). Within the BOLD activation mask, 77.2% of voxels showed no statistically significant group-level ΔCMRO_2_ effect and therefore could not be robustly classified as concordant or discordant. Among the remaining voxels (22.8%), 17.4% of all voxels were classified as concordant and 5.4% as discordant (Fig. 3A–B).

**Figure 3.**
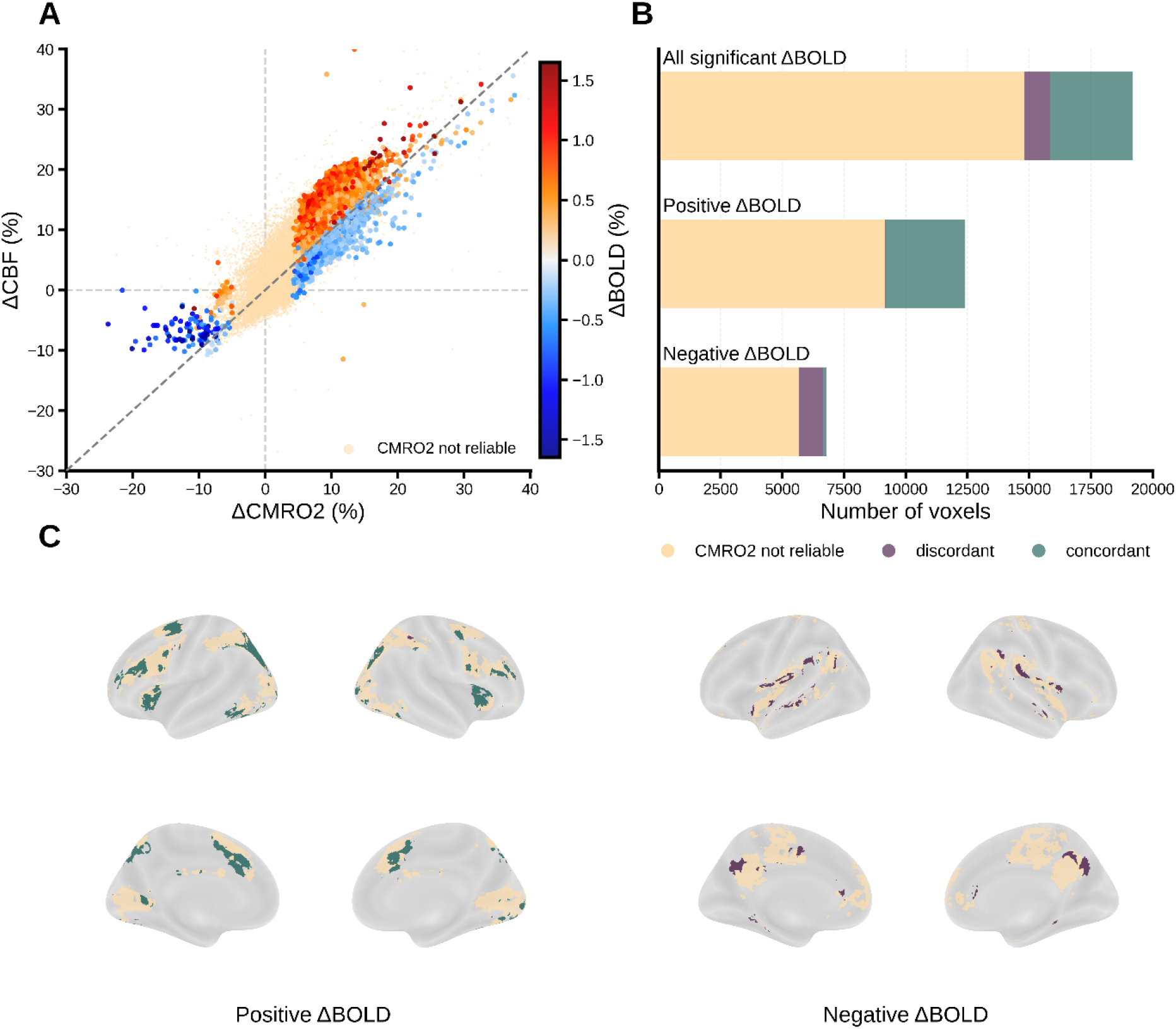
**A** Relationship between median task-evoked changes in cerebral blood flow (ΔCBF) and cerebral metabolic rate of oxygen (ΔCMRO_2_) across all voxels showing statistically significant ΔBOLD responses. Voxels with statistically reliable ΔCMRO_2_ are color-coded by ΔBOLD sign and magnitude, whereas voxels with non-significant ΔCMRO_2_ are shown in light orange. The dashed diagonal indicates equal relative changes in CBF and CMRO_2_; voxels above the diagonal for negative ΔCMRO_2_ and below the diagonal for positive ΔCMRO_2_ correspond to an n-ratio (ΔCBF/ΔCMRO_2_) < 1. **B** Distribution of voxels within the BOLD activation mask by CMRO_2_ reliability and sign concordance. The top bar shows all voxels with significant ΔBOLD responses (n = 19,190), partitioned into voxels with no significant ΔCMRO_2_ effect (77.2%), concordant voxels (17.4%), and discordant voxels (5.4%). The middle bar shows voxels with positive ΔBOLD, of which 26.0% were classified as concordant, 0.4% as discordant, and 73.6% showed no significant ΔCMRO_2_ effect. The bottom bar shows voxels with negative ΔBOLD, of which 1.7% were concordant, 14.6% discordant, and 83.7% showed no significant ΔCMRO_2_ effect. Bar lengths indicate the number of voxels in each category. **C** Cortical surface maps illustrating the spatial distribution of voxel classifications shown in (B). Left panels display voxels with positive ΔBOLD; right panels display voxels with negative ΔBOLD. Colors correspond to concordant, discordant, and CMRO_2_ not reliable classifications.

Clear differences in sign concordance patterns were observed between positive and negative BOLD activation maps (Fig. 3B,C). Within the positive BOLD mask, discordance was rare (0.4%), whereas 26% of voxels were concordant and 73.6% showed no statistically significant ΔCMRO_2_. In contrast, within the negative BOLD mask, 14.6% of voxels were discordant and 1.7% concordant, while 83.7% showed no significant ΔCMRO_2_ effect. Crucially, these results indicate that, for most voxels classified as concordant or discordant by Epp et al., ΔCMRO_2_ estimates do not provide sufficient statistical support for robust sign determination. Where classification was possible, positive ΔBOLD was overwhelmingly concordant with metabolism, whereas negative ΔBOLD exhibited considerable higher rates of sign opposition, consistent with prior PET–fMRI findings (Godbersen et al., 2023; Stiernman et al., 2021).

Taken together, our analyses indicate that the central claim of Epp et al. – that approximately 40% of voxels exhibit metabolic responses opposite in sign to the observed BOLD signal – is not supported once the statistical robustness of ΔCMRO_2_ estimates is taken into account. Rather than indicating systematic physiological sign reversal, the reported voxel-wise discordance appears largely attributable to substantial variability in ΔCMRO_2_ estimates across participants, with 77.2% of voxels showing no statistically reliable direction of metabolic change. While some variability may reflect genuine inter-individual differences, it likely also reflects elevated noise levels for quantitative fMRI-based CMRO_2_ estimation. When this variability is taken into account, the resulting pattern is largely consistent with prior literature: negative BOLD responses remain difficult to interpret due to heterogeneous underlying mechanisms, whereas for positive BOLD responses, where ΔCMRO_2_ effects reached statistical significance at the group level, metabolic changes were predominantly concordant with the observed BOLD signal. We therefore view the study by Epp et al. as an important contribution that stimulates discussion about BOLD interpretation, while suggesting that its conclusions should be considered in light of the statistical uncertainty of the underlying metabolic estimates.

Finally, we would like to commend Epp et al. for their commitment to open science principles. By making their data publicly available, they have enabled the kind of critical dialogue that is essential for scientific progress, rigor and transparency of the field.

## Materials and Methods

### Dataset and Participants

We reanalysed the openly available dataset used by Epp et al. (2025). Data were retrieved from OpenNeuro (https://openneuro.org/datasets/ds004873). Of the 40 participants included in the original study, we were able to retrieve complete data for 38 participants; two participants were excluded due to missing derivatives. All analyses were conducted on these 38 participants.

MRI data were acquired on a 3 T Philips Ingenia MRI scanner equipped with a 32-channel head coil. The quantitative fMRI protocol combined multiparametric quantitative BOLD (mqBOLD) imaging with arterial spin labeling (ASL) to estimate hemodynamic and metabolic parameters. Multiecho spin-echo T2 mapping was acquired using a 3D gradient spin-echo readout with eight echoes (TE_1_ = ΔTE = 16 ms; TR = 251 ms; flip angle = 90°; voxel size = 2 × 2 × 3.3 mm^3^; 35 slices). Multiecho gradient-echo T2* mapping was acquired with 12 echoes (TE_1_ = ΔTE = 5 ms; TR = 2,229 ms; flip angle = 30°; voxel size = 2 × 2 × 3 mm^3^; 35 slices). Cerebral blood volume (CBV) was measured using dynamic susceptibility contrast (DSC) MRI with single-shot GRE-EPI (TR = 2.0 s; flip angle = 60°; voxel size = 2 × 2 × 3.5 mm^3^; 35 slices; 80 dynamics) following administration of a gadolinium-based contrast agent (0.1 ml kg^−1^). Cerebral blood flow (CBF) was measured using pseudocontinuous arterial spin labeling (pCASL; post-labeling delay = 1,800 ms; labeling duration = 1,800 ms; TE = 11 ms; TR = 4,500 ms; voxel size = 3.28 × 3.5 × 6.0 mm^3^; 20 slices; 39 dynamics plus M0 scan). Task-based BOLD fMRI was acquired using single-shot echo-planar imaging (TR = 1.2 s; TE = 30 ms; flip angle = 70°; voxel size = 3 × 3 × 3 mm^3^; 40 slices; multiband factor = 2; 400 dynamics). A B0 field map was acquired for susceptibility correction (TR = 525 ms; TE_1_/TE_2_ = 6.0/9.8 ms; voxel size = 3 × 3 × 3 mm^3^). These measurements were combined in the original study to derive voxel-wise maps of oxygen extraction fraction (OEF) and cerebral metabolic rate of oxygen consumption (CMRO_2_) using the mqBOLD framework and Fick’s principle. Full acquisition details are reported in Epp et al. (2025).

### Task

Participants performed four task conditions in the original experiment: a calculation task (CALC), an autobiographical memory task, a low-level control condition (CTRL), and a resting-state baseline, as described in Epp et al. (2025). In the present reanalysis we focused exclusively on the CALC and CTRL conditions, as these were the main focus of Epp et al. as well.

In the CALC condition, participants solved arithmetic problems presented visually. Each trial displayed a row of three numbers followed by a question mark (n_1_, n_2_, n_3_, ?), and participants were instructed to determine the missing number according to the arithmetic rule governing the sequence. Responses were selected from three answer options using the button box, with a maximum response time of 10 s per trial.

In the CTRL condition, participants performed a low-level baseline task with minimal cognitive demand. A row of random letters was presented for several seconds, and participants indicated via button press whether the first letter was a vowel. This condition was intended to provide visual input and motor responses comparable to the CALC task.

### Preprocessing and Spatial Normalization

We did not perform preprocessing of the raw MRI data. Instead, we used the derivative datasets released by Epp et al. together with the original publication (https://openneuro.org/datasets/ds004873). The full preprocessing and modelling pipeline is described in detail in Epp et al. (2025); here we briefly summarize the relevant steps.

Preprocessing of BOLD fMRI data included motion correction, susceptibility distortion correction using field maps, as well as estimation and removal of confound regressors. Functional images were coregistered to the individual T1-weighted anatomical image using boundary-based registration implemented in FSL, and subsequently normalized to MNI152NLin6Asym standard space (2 mm isotropic resolution) using nonlinear registration implemented in ANTs 2.3.3.

Quantitative parameter maps used for metabolic modelling were computed in subject space from multiparametric qBOLD and ASL data using in-house scripts (MATLAB) and SPM12, as described by Epp et al. These included maps of T2, T2*, R2′, cerebral blood flow (CBF), cerebral blood volume (CBV), oxygen extraction fraction (OEF), and cerebral metabolic rate of oxygen consumption (CMRO_2_). All parameter maps were first coregistered to the first echo of the T2 acquisition and subsequently aligned to the participant’s native T1-weighted anatomical image.

Spatial normalization to standard space was performed by applying the T1 to MNI nonlinear transformation estimated by fMRIPrep to all parameter maps. Where MNI-space derivatives were not directly available in the shared dataset, native-space parameter maps were transformed into MNI152 space using the transformation fields provided by the original authors.

All analyses were conducted in MNI152 space at 2 mm isotropic resolution using the group-level gray matter mask provided with the derivative dataset. To ensure consistency with the original study, we exclusively used the transformation fields and processing scripts released by Epp et al. (https://github.com/NeuroenergeticsLab/two_modes_of_hemodynamics) and did not introduce additional spatial preprocessing steps.

### Percent Signal Change Maps

To ensure consistency with the original analysis, we used the percent signal change (PSC) maps provided in the derivative dataset released by Epp et al. (2025). These maps quantify task-evoked BOLD signal changes for the CALC condition relative to the CTRL baseline and are provided in MNI152 standard space (2 mm isotropic resolution).

PSC maps were derived from the preprocessed BOLD time series using the procedure implemented in the original preprocessing pipeline. For each participant and voxel, percent signal change was calculated as the relative difference between the BOLD signal during the CALC task and the baseline signal during the CTRL condition, expressed as a percentage of the baseline signal. Voxels with invalid baseline values were excluded during PSC computation to avoid numerical instability.

Task-evoked changes in CMRO_2_ were computed by comparing CMRO_2_ estimates obtained for the CALC and CTRL conditions. In the original study, CMRO_2_ maps were derived using a semiquantitative, BOLD-informed approach in which baseline quantitative parameter maps (R2′, CBV, and CBF) were combined with task-related BOLD signal changes to estimate condition-specific CMRO_2_ via the mqBOLD framework and Fick’s principle. In this approach, task-related changes in R2′ are approximated from baseline R2′ values and the observed BOLD signal change, rather than being estimated directly from multiecho acquisitions during task conditions, with the aim of reducing noise propagation. Percent signal change maps for CMRO_2_ were then calculated as the relative difference between CALC and CTRL CMRO_2_ estimates.

### Voxel Selection: BOLD Activation Mask

Similar to Epp et al. (2025), we restricted subsequent analyses to voxels showing statistically significant task-evoked BOLD responses at the group level. In the original study, significant task-related BOLD effects were identified using a multivariate partial least squares (PLS) analysis that extracted latent variables relating BOLD activity to task conditions. In the present analysis, we instead defined the BOLD activation mask using a voxel-wise univariate approach to enable straightforward statistical interpretation. In contrast to voxel-wise statistical testing, PLS evaluates significance at the level of latent variables, while voxel contributions are interpreted using bootstrap ratios rather than formal voxel-wise p-values. Consequently, thresholded bootstrap maps may contain large numbers of voxels exceeding the chosen threshold without providing direct voxel-wise significance inference. We therefore adopted a conventional voxel-wise statistical framework to define task-responsive voxels. For each voxel, a one-sample t-test against zero was performed across participants on the CALC–CTRL percent signal change values. Resulting p-values were corrected for multiple comparisons using Benjamini–Hochberg false discovery rate (FDR) correction at q = 0.05. Voxels surviving correction defined the BOLD activation mask. Positive and negative ΔBOLD voxels were considered separately in subsequent analyses where indicated.

### Coefficient of Variation Analysis

To quantify inter-individual variability in measured parameters, we computed voxel-wise coefficients of variation (CV) across participants. The CV was defined as the ratio of the standard deviation to the absolute value of the mean across participants (CV = SD/|mean|) and was calculated separately for each voxel in MNI space.

CV values were computed for baseline (CTRL) and task (CALC) conditions using the corresponding parameter maps provided in the derivative dataset. Specifically, CV distributions were evaluated for CBF, CBV, T2*, R2′, and CMRO_2_. For each parameter and condition, voxel-wise CV values were calculated across participants and subsequently aggregated across cortical voxels to obtain distributional summaries.

In addition, CV values were computed for PSC maps derived from the CALC–CTRL contrast. PSC maps were available for BOLD and the quantitative parameters used in the mqBOLD framework, allowing direct comparison of relative variability in task-evoked signal changes across modalities.

For visualization, voxel-wise CV values were pooled across voxels and plotted as kernel density distributions for each parameter. Separate distributions were generated for baseline (CTRL), task (CALC), and PSC maps (Fig. 1). These distributions were used to compare the relative variability of physiological parameters across participants under baseline conditions, during task performance, and for task-evoked signal changes.

As CV depends on the magnitude of the mean, CV values may be inflated in regions where mean signal changes are close to zero. The comparison therefore primarily serves as a descriptive indicator of relative variability rather than a formal statistical test.

### Voxel-wise Sign Discordance (Replication Analysis)

To replicate the primary analysis of Epp et al., we computed voxel-wise sign discordance between group-mean ΔBOLD and group-mean ΔCMRO_2_ within the BOLD activation mask. Voxels were classified as concordant when the signs of the group-averaged ΔBOLD and ΔCMRO_2_ matched, and as discordant when the two signals exhibited opposing signs.

### Participant-level Sign Concordance Analysis

To assess the robustness of voxel-wise sign classification across participants, we computed, for each voxel, the proportion of participants showing concordant sign between ΔBOLD and ΔCMRO_2_. A concordance rate of 0.5 indicates equal concordance and discordance across participants. We tested whether the observed concordance proportion differed significantly from 0.5 using a binomial test at each voxel. Resulting p-values were corrected for multiple comparisons using FDR (q = 0.05). Voxels were classified as:

1. Concordant: significantly greater concordance than 0.5,
2. Discordant: significantly less concordance than 0.5,
3. Indeterminate: not significantly different from 0.5.

### Group-level ΔCMRO_2_ Classification

As a complementary and typically less conservative approach, we evaluated whether ΔCMRO_2_ differed significantly from zero at the group level using a one-sample t-test across participants. Resulting p-values were corrected for multiple comparisons using FDR (q = 0.05). This approach is typically less conservative than the participant-level concordance test because it evaluates whether the group mean differs from zero and does not require a supermajority of participants to exhibit the same sign relationship.

Within the BOLD activation mask, voxels were categorized as follows:

1. Concordant: ΔCMRO_2_ significantly different from zero and matching the sign of ΔBOLD,
2. Discordant: ΔCMRO_2_ significantly different from zero and opposite in sign to ΔBOLD,
3. No significant ΔCMRO_2_: ΔCMRO_2_ not significantly different from zero.

This classification allowed us to determine in how many voxels the direction of metabolic change could be inferred with statistical support.

### Statistical Thresholds and Software

All statistical analyses were conducted in Python using NumPy, SciPy and Statsmodels. One-sample t-tests (scipy.stats.ttest_1samp) were used for group-level mean testing. Binomial tests were used for participant-level sign consistency. Multiple comparisons were controlled using Benjamini–Hochberg FDR correction with q = 0.05. Figures were generated using Matplotlib and Nilearn for surface visualization.

## Data Availability

The original dataset generated by Epp et al. is publicly available via OpenNeuro (https://openneuro.org/datasets/ds004873/versions/2.0.6). All scripts used for data processing, statistical analyses, and figure generation, as well as summary statistics associated with this manuscript, are available in a public GitHub repo: https://github.com/olegolt/BOLD_metabolism_reanalysis

